# Global analysis of plasma lipids identifies liver-derived acyl-carnitines as a fuel source for brown fat thermogenesis

**DOI:** 10.1101/132241

**Authors:** Judith Simcox, Gisela Geoghegan, John Alan Maschek, Amanda Mixon, Marzia Pasquali, Ren Miao, Sanghoon Lee, Lei Jiang, Ian Huck, Anthony J. Donato, Udayan Apte, Nicola Longo, Jared Rutter, James Cox, Claudio J. Villanueva

## Abstract

Cold induced thermogenesis is an energy demanding process that protects endotherms against a reduction in ambient temperature. Using non-targeted LC-MS based lipidomics, we identified plasma acylcarnitines as the most significantly changed lipid class in response to the cold. Here we show that acylcarnitines provide fuel for brown fat thermogenesis. In response to the cold, FFAs released from adipocytes activate the nuclear receptor HNF4α to stimulate the expression of genes involved in acylcarnitine metabolism in the liver. Conditional deletion of HNF4α in hepatocytes blocks the cold-induced changes in hepatic gene expression, lowering circulating long chain acylcarnitine (LCAC) levels, and impairing their ability to adapt to the cold. Finally, a bolus of L-carnitine or palmitoylcarnitine rescues the cold sensitivity seen with aging. Our data highlights an elegant mechanism whereby white adipose tissue provides FFAs for hepatic carnitilation to generate plasma LCAC as a fuel source for BAT thermogenesis.

**Highlights:**

- Blood acylcarnitine levels increase in response to the cold.
- FFA mobilization in response to the cold activates hepatic HNF4α and stimulates genes involved in acylcarnitine metabolism.
- Brown adipocytes metabolize palmitoylcarnitine.
- Carnitine administration improves thermogenic response in aged mice.

**ETOC:** Simcox et al identified acylcarnitines as a novel source of energy for thermogenesis. In response to the cold, the liver activates a transcriptional program through the transcription factor HNF4α, leading to increased acylcarnitine levels. They also find that aging mice have reduced acylcarnitine levels and an impaired thermogenic response in the cold. Increasing acylcarnitine levels in old mice increases their ability to adapt to the cold. Their studies discover a physiological role for acylcarnitines in thermogenesis.

**Graphical Abstract:** Cold exposure stimulates the sympathetic nervous system to release noradrenaline (NA). Activation of β3-adrenergic receptors stimulates FFA release and activation of the transcription factor HNF4α in the liver. This leads to increased gene expression of enzymes involved in acylcarnitine metabolism. The acylcarnitines are released in the blood to provide fuel for brown fat thermogenesis. These studies highlight the role of the liver in the thermogenic response.

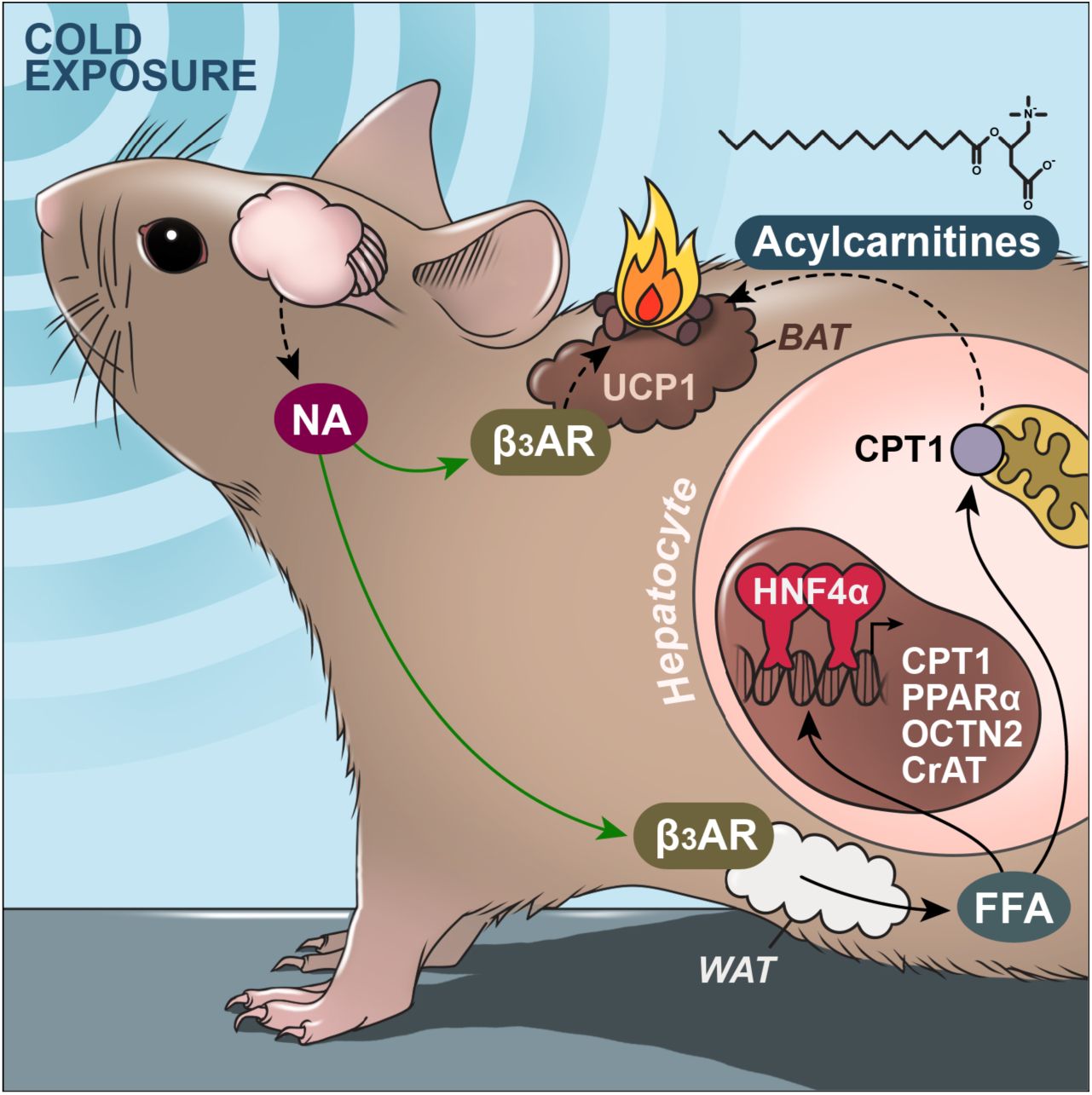

## Introduction

Cellular metabolic networks evolved through the selective pressures of starvation and cold exposure. In mammals, these two energetically challenging conditions require dynamic communication between tissues to maintain survival. During starvation, catecholamines signal to β-adrenergic receptors in the white adipose tissue to mobilize lipid stores as free fatty acids. Preferential use of circulating fatty acids by the liver and muscle during fasting preserves glucose stores for the brain. Throughout this metabolic switch the liver uniquely provides ketones that can be metabolized by neurons in the Central Nervous System (CNS) to generate ATP. This switch in primary fuel sources between tissues is regulated by metabolic substrate competition and inhibitory signaling in the Randle cycle (Hue and Taegtmeyer, 2009). Similar pathways are upregulated during cold exposure where β3-adrenergic receptor activation leads to increased triglyceride lipolysis in white adipocytes, and activation of thermogenesis in brown adipocytes (Lafontan and Berlan, 1993). However, little is known about the metabolic adaptations that occur in other cell types during acute cold challenge.

The brown adipose tissue (BAT) plays a major role in protecting against the cold through non-shivering thermogenesis. Brown adipocytes generate heat by disrupting ATP synthesis in the mitochondria, allowing protons to leak out through the uncoupling protein 1 (UCP1), releasing the potential energy as heat. Mice lacking UCP1 develop hypothermia with acute cold challenge, however these mice are able to adapt with incremental reduction in ambient temperature, suggesting that alternative mechanisms of thermogenesis are at play (Golozoubova et al., 2001). To replenish the proton gradient, brown adipocytes increase utilization of both glucose and fatty acids, generating additional heat as a byproduct of cellular metabolism (Seale et al., 2009). During cold exposure the BAT relies on glucose, and when activated increases glucose uptake by 12-fold (Orava et al., 2011; Yu et al., 2002). In addition to carbohydrates, brown adipocytes utilize fatty acids from triglyceride rich lipoproteins and free fatty acids released by white adipocytes (Bartelt et al., 2011; Chondronikola et al., 2014). Mice lacking adipose triglyceride lipase (ATGL) in adipocytes are unable to maintain their body temperature during a cold challenge, highlighting the importance of energy mobilization for thermogenesis (Haemmerle et al., 2006). However, beyond glucose and free fatty acids, there is little known about additional fuel sources that drive thermogenesis.

Mitochondrial fatty acid oxidation in the liver is a tightly regulated process that is activated upon fasting. After entry into cells, long-chain fatty acids are activated by acyl-CoA synthetase, and conjugated with carnitine by CPT1, the rate limiting enzyme in long-chain fatty acid oxidation (Esser et al., 1993; Fingerhut et al., 2001; Longo et al., 2006; Schooneman et al., 2013). Carnitine is transported into cells through a cell surface transporter, Octn2 (Tamai et al., 1998). Conjugation with carnitine allows the transport of fatty acids across the inner mitochondrial membrane through the carnitine-acylcarnitine translocase (CACT) (Ramsay et al., 2001). The carnitine group is then removed, and fatty acids are destined for oxidation after activation by CPT2 to generate fatty acyl-CoAs (Gempel et al., 2002). CPT1 is regulated by the transcription factor HNF4α, a nuclear receptor that plays a key role in liver development and mitochondrial energetics (Martinez-Jimenez et al., 2010). Mutations in HNF4α lead to maturity-onset diabetes of the young type 1 (MODY 1), a disorder characterized by defective glucose-stimulated insulin secretion (Yamagata et al., 1996). During exercise and fasting acyl-carnitines are elevated in the plasma, and are thought to reflect incomplete fatty acid oxidation (Schooneman et al., 2013). Acyl-carnitines are also elevated in several inborn errors of metabolism, including disorders of fatty acid oxidation where mutations in MCAD, VLCAD, and LCHAD can lead to death (Shekhawat et al., 2005). Although they have been identified in the plasma, little is known about their role in systemic energy metabolism, and their regulation.

In this study, we performed non-targeted lipidomics of plasma, and found that long chain acylcarnitines were induced in response to cold exposure or treatment with β3 adrenergic receptor agonist CL-316,243. We hypothesized that acylcarnitines served a greater role than being a byproduct of fatty acid oxidation, but rather a mechanism to provide a fuel source for BAT thermogenesis. These studies led to a previously unappreciated role for hepatic HNF4α in regulating cold-induced changes in expression of enzymes involved in hepatic acylcarnitine metabolism. We demonstrate that this transcriptional program requires ATGL-mediated lipolysis of FFAs to activate HNF4α. With aging, mice show reduced plasma acylcarnitine levels in response to the cold, and display a cold sensitive phenotype. This can be reversed with carnitine or palmitoyl-carnitine supplementation. Our findings suggest a novel role for the liver in thermogenesis, as well as uncovering a well-orchestrated inter-tissue communication system to upregulate energy mobilization for heat production.

## Results

### Global lipid analysis of plasma from mice exposed to cold and the identification of acylcarnitines

To understand the metabolic changes that occur during acute cold exposure, mice were exposed to room temperature (24°C) or cold (4°C) for 5 hours, and plasma was analyzed using LC-MS based lipidomic analysis, and lipids were displayed as a heatmap after cluster analysis using MetaboAnalyst 3.0 (Figure 1A). A total of 287 circulating lipids significantly changed (p≤0.05) in response to cold exposure, and 93 lipids were elevated, while 194 lipids were down-regulated. Further analysis and identification of the lipids or derivatives that were significantly (p<0.01) upregulated by two fold showed a high representation of long-chain acylcarnitine (LCAC) species (Figure 1B). We confirmed that these lipids were LCACs using tandem mass spectroscopy (LC-MS/MS). To determine if the elevated LCACs reflected a change in thermogenic potential, we utilized aged mice as a physiologically relevant model of impaired thermogenesis. As mice and humans age, there is a loss of BAT function and increased sensitivity to hypothermia (Sellayah and Sikder, 2014; Yoneshiro et al., 2011). Therefore, we compared plasma lipidomic profiles between 3 month-old mice and 24-month old mice, and found a temperature dependent divergence in the plasma lipid signature (Figure 1C), where older mice have a blunted response in the cold. To test whether LCAC species were reduced in older mice, we looked at 14:0-Carnitine, 16:1-Carnitine, 18:0-Carnitine, and 18:2-Carnitine levels in the plasma, and found that 14:0-Carnitine was elevated twelve-fold in response to the cold, compared to a modest increase of two-fold in 24month old mice (Figure 1D). In our initial analysis we only identified long chain acylcarnitines, and we were curious whether other acylcarnitine species were altered. Using ultra performance LC-MS/MS (UPLC-MS/MS) we quantified many of the previously identified acylcarnitine species and found that the sum of short-chain acylcarnitines (SCAC) (C2-C8), medium-chain acylcarnitines (MCAC) (C10-C14), and long-chain acyl carnitines (LCAC) (C16-C18) (Figure 1E, **FigureS1**) were elevated in response to cold exposure, with the greatest changes occurring in 3 month-old mice. Notably, older mice had higher basal levels of LCAC, MCAC, and SCAC at room temperature, but a clear blunted response in the cold (**Figure S2A**). In contrast, plasma carnitine levels were reduced in response to the cold in both 3-month and 24-month old mice (Figure 1E). These results are consistent with impaired thermogenesis seen in 24-month old mice (**Figure S2A**), and suggest a loss of metabolic flexibility that impairs the response to energetically stressful stimulus.

**Figure 1:**
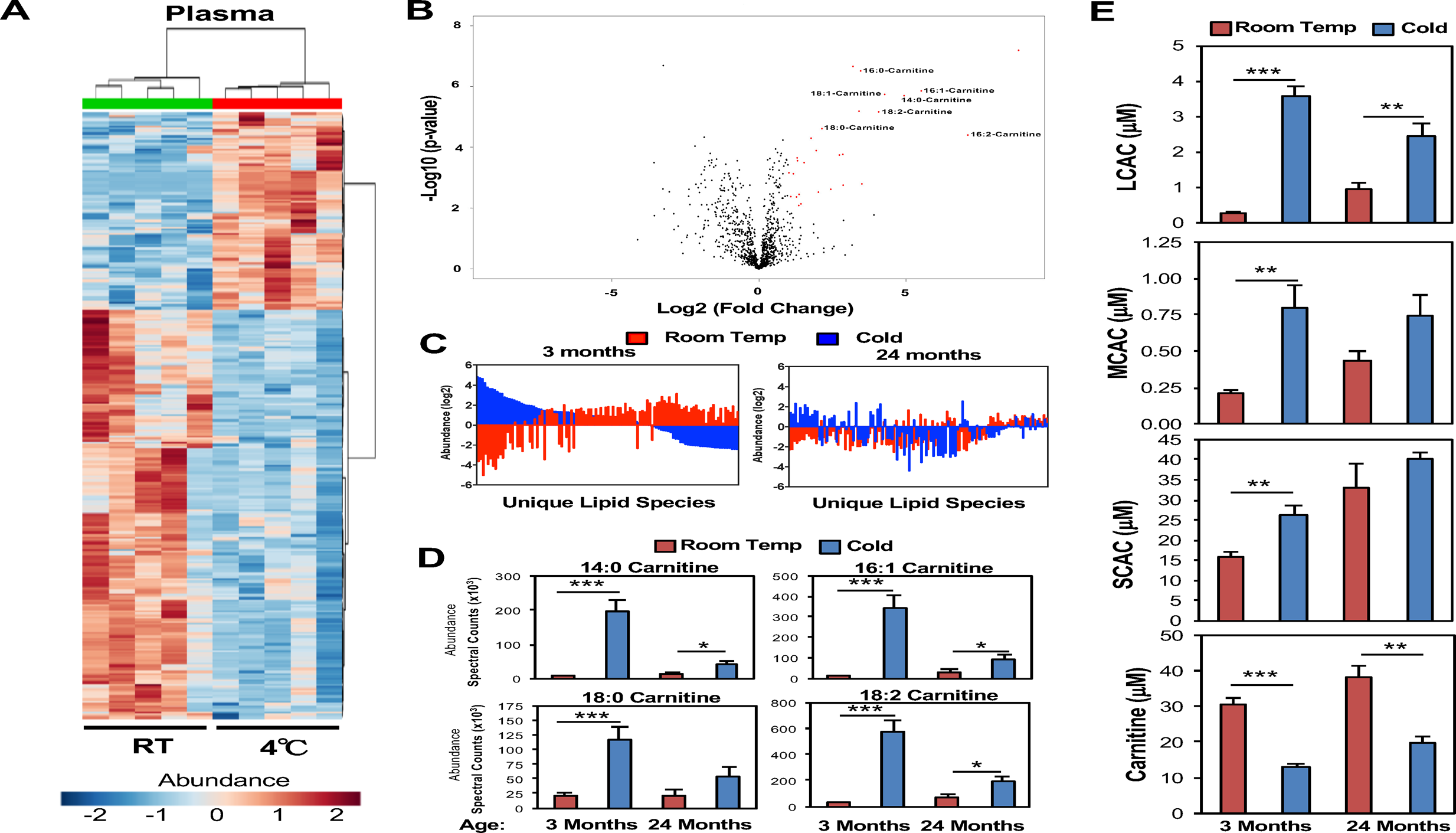
Acylcarnitine levels in the blood are elevated in response to cold exposure. (A) Heat Map and cluster analysis of 281 lipids from plasma of 3-month old C57BL6J male mice at room temperature (RT) or Cold (4°C). Samples were run on an Agilent 6490 LC-MS and analyzed using Metaboanalyst 3.0 (N = 5/group). (B) Volcano plot of LC-MS based lipidomics from the plasma of 3-month old C57BL6J male mice at room temperature (RT) or Cold (4°C). Lipids that are increased 2-fold in Cold/RT and have a p-value below 0.01 are in labeled in red. Long chain acylcarnitine species were identified through LC-MS/MS (N = 5/group). (C) SAM analysis of LC-MS lipidomics of plasma from room temperature vs cold exposed in C57BL6J male mice 3-month or 24-month old (N=4-5/group). (D) Plasma acylcarnitine levels of 3-month and 24-month old C57BL6J male mice at room temperature (RT) or Cold (4°C) (N=4-5/group). (E) Plasma levels of long chain acylcarnitine (LCAC), medium chain acylcarnitines (MCAC), short chain acylcarnitines (SCAC), and carnitine in the plasma of 3 month and 24 month old C57BL6J male mice as measured by LC-MS (N=4-5/group). Data are presented as mean ± SEM. *p ≤ 0.05, **p ≤ 0.01, ***p≤0.001. See also Figure S1.

### Expression of genes involved in acylcarnitine metabolism induced by cold exposure and β3 adrenergic receptor agonist

Circulating acylcarnitine levels change during acute energetic stress including exercise and fasting (Costa et al., 1999; Hiatt et al., 1989; Yamaguti et al., 1996). These acute changes are regulated in part by transcriptional changes in components of the carnitine shuttle (Song et al., 2010; Vila-Brau et al., 2013). To test which tissues were contributing to the increase in acylcarnitine levels, we measured the expression of genes involved in acylcarnitine metabolism in the liver, skeletal muscle, and BAT. In the liver we found the expression of Cpt1b, Octn2, Cact, Cpt2, and Crat were increased in response to cold exposure, while the expression of Bbox1, an enzyme involved in carnitine synthesis, was not changed (Figure 2A). These changes were also reflected in the liver by western blot analysis using antibodies that detect Cpt1b and Octn2 (Figure 2B). In contrast, there were no detectable changes in expression of Cpt1b, Octn2, CrAT, and CACT in the skeletal muscle (**Figure S2C**), or BAT, except for increased expression of Octn2 (Figure 2C). To test whether expression of these genes mirrored the blunted response in acylcarnitines seen with aging, we measured the expression of Cpt1b, Octn2, CrAT, CACT, and found that the induction with cold exposure was lost in the liver of the 24 month old mice, correlating with the lower circulating LCAC and impaired thermogenesis seen in 24 month old mice (Figure 2D, Figure 1D, **Figure S2A**). These data suggest that transcriptional changes in hepatic acylcarnitine metabolism contribute to the blunted induction in plasma acylcarnitine levels seen with aging. To test whether the increase in circulating acylcarnitines was due to brown adipose tissue, we measured LCACs in UCP1-DTA mice that lack BAT. UCP1-DTA mice were previously generated by expressing diphtheria toxin using the UCP1 promoter (Lowell et al., 1993). In response to the cold, UCP1-DTA mice have a similar induction in LCACs when compared to littermate controls (**Figure S2D**). Both 14:0-Carnitine and 18:1-Carnitine in the blood were elevated approximately 2 fold in response to a 4hr cold exposure. Although the basal levels trend to be lower, the similar fold induction in control and mice lacking BAT, suggests that the source of acylcarnitines is not brown adipocytes.

**Figure 2:**
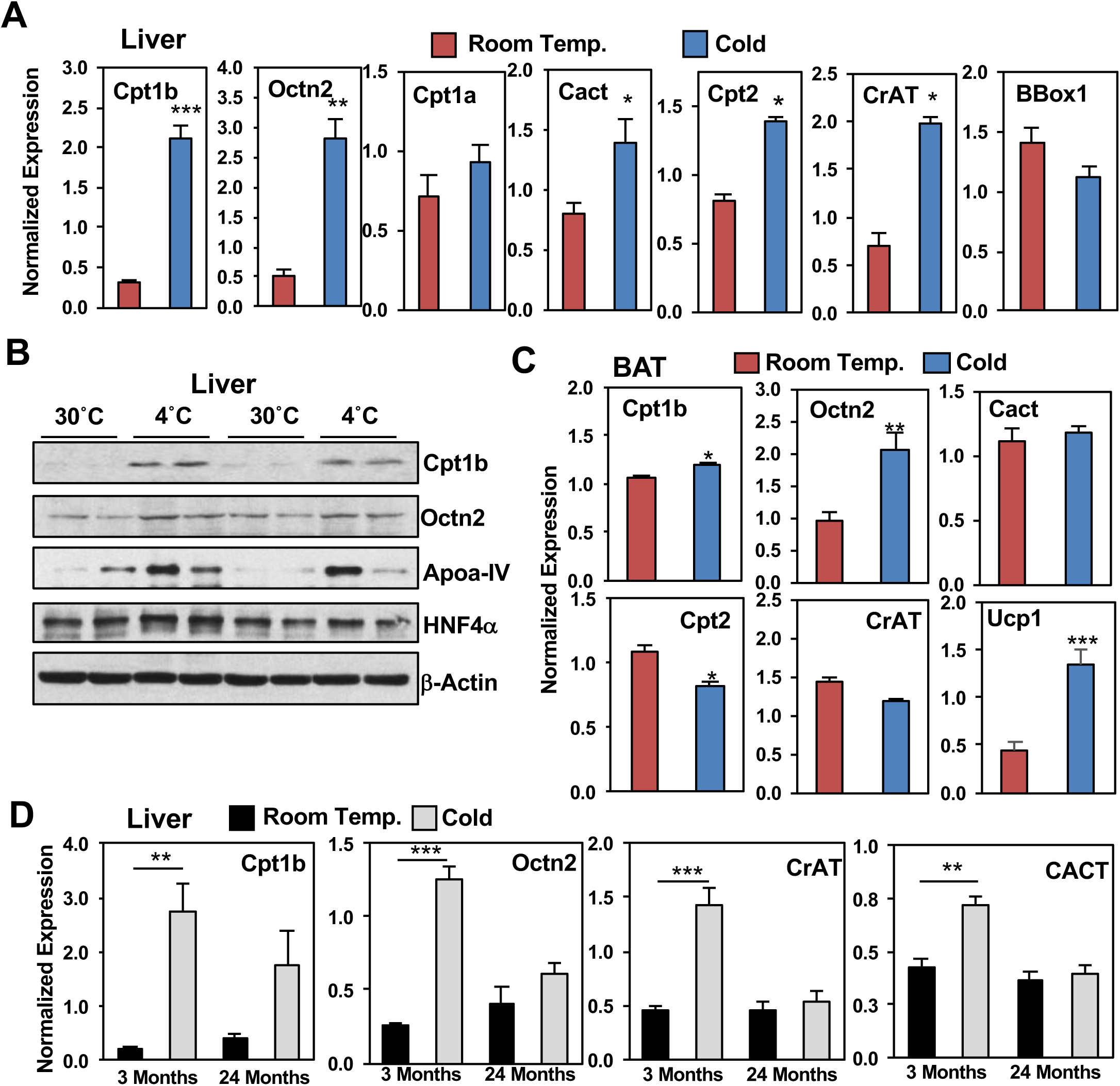
Genes involved in acylcarnitine transport and metabolism are increased in the liver of cold exposed mice. (A) Gene expression in livers of 3-month old C57BL6J male mice at room temperature (RT) or Cold (4°C) for 5hrs. N = 5/group. (B) Western blot analysis of livers detecting CPT1b, HNF4α, and HMGB1 in 3-month old C57BL6J male mice at thermoneutrality (30°C) or Cold (4°C) for 5hrs. N = 5/group. (C) Gene expression of skeletal muscle (gastrocnemius) of 3-month old C57BL6J male mice at room temperature (RT) or Cold (4°C) for 5hrs. N = 5/group. (D) Gene expression of brown adipose tissue (BAT) of 3-month old C57BL6J male mice at room temperature (RT) or Cold (4°C) for 5hrs. N = 5/group. (E) Gene expression in livers of 3-month and 24-month old C57BL6J male mice at room temperature (RT) or Cold (4°C) for 5hrs. N = 5/group. All transcripts were normalized to RPL13. Data are presented as mean ± SEM. *p ≤ 0.05, **p ≤ 0.01, ***p≤0.001. See also Figure S2.

During cold exposure body temperatures are maintained through both shivering and non-shivering thermogenesis. There is evidence that exercise leads to elevated acylcarnitine levels, so we hypothesized that shivering might trigger an induction in acylcarnitines as well. To rule that out, we used a stimulus of thermogenesis where shivering is not activated. Activation of non-shivering thermogenesis is regulated by the sympathetic nervous system through activation of β3 adrenergic receptors (β3AR), which are found in both white and brown adipocytes. Mice were treated with the β3-AR agonist CL-316,243 (Himms-Hagen et al., 1994) or vehicle control, and we found that CL-316,243 stimulates serum acylcarnitines, as noted by the elevated levels of 12:0-,14:0-,14:1-, 16:0-, 16:1-, 16:2-, 18:0-, 18:1-, and 18:2-Carnitine (Figure 3A). To test whether CL-316,243 stimulates hepatic gene expression of enzymes involved in acylcarnitine metabolism as well, we measured Cpt1b, Octn2, Cpt1a, Cact, Cpt2, and Crat mRNA, and found an induction in the liver (Figure 3B), with little effects seen in BAT (**Figure S3A**). However, we did not see changes in expression of genes involved in lipid mobilization and metabolism, including fatty acid translocase (CD36) and acetyl-CoA acetyltransferase (Acat1) (Figure 3C). In response to CL-316,243 administration, a subset of BAT acylcarnitines increased, however acylcarnitine levels in the liver were largely unchanged (Figure S3B and S3C). CL-316,243 is a selective β3-AR agonist, yet it was able to stimulate changes in hepatic gene expression despite the lack of expression of the β3-AR in the liver. This led us to test whether other external stimuli could activate hepatic gene expression, in particular FFA mobilization from adipocytes. Hepatocytes respond to FFAs through the HNF4α transcription factor, of which several direct targets are known, including MTTP, ApoA-IV, PPARα, all of which were upregulated in response to CL-316,243 administration and cold exposure (Figure 3C and Figure 2B) (Hanniman et al., 2006; Martinez-Jimenez et al., 2010; Sheena et al., 2005) However, not all HNF4α targets are induced, both ApoC3 and Acadm expression were unaltered in response to CL-316,243 administration, suggesting a unique transcriptional response.

**Figure 3:**
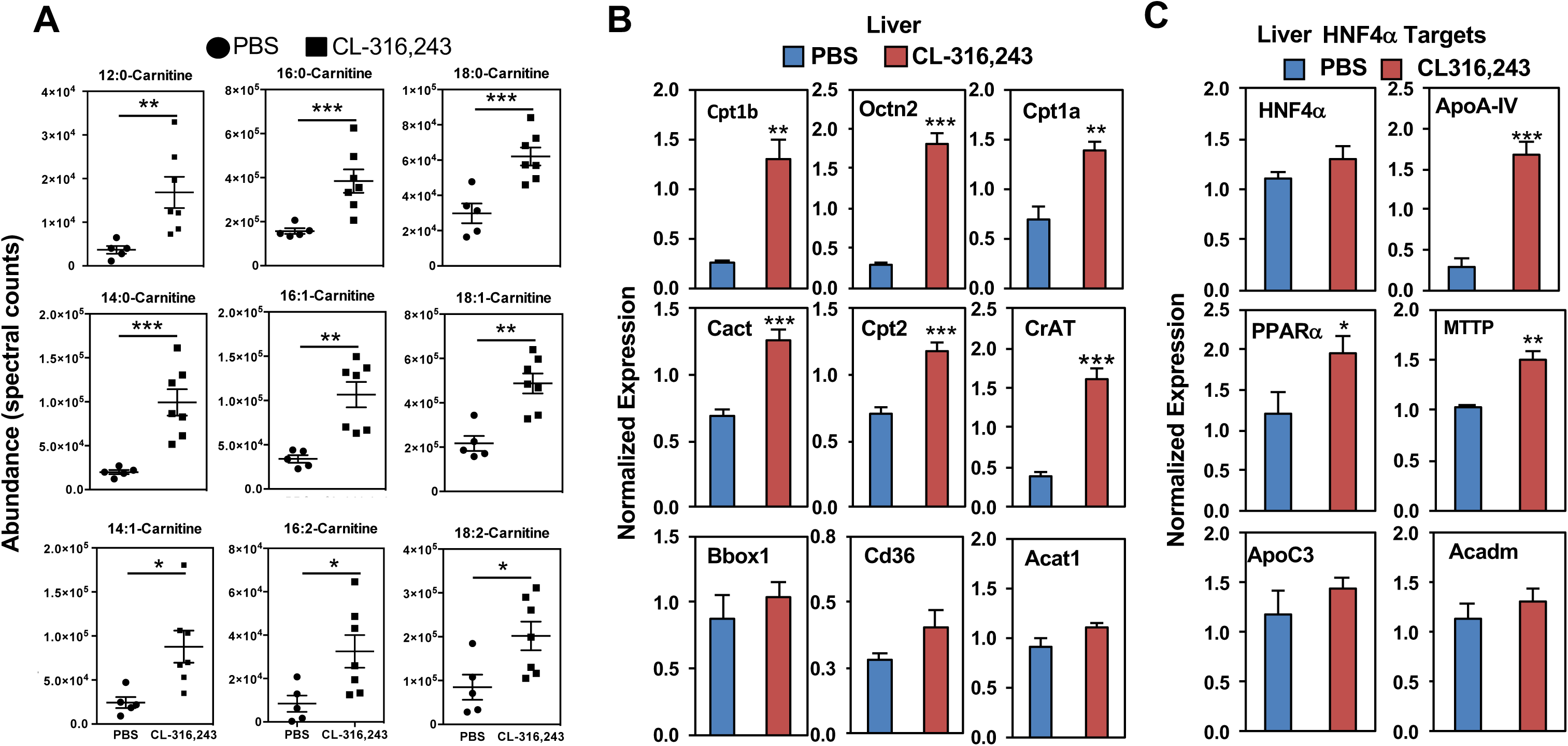
β3AR agonist CL-316,243 increases blood acylcarnitine levels and stimulates hepatic transcripts involved in acylcarnitine synthesis. (A) Serum acylcarnitine levels as measured by LC-MS in 3-month old mice treated with PBS or CL-316-243. N = 5-7/group. (B) Expression of gene involved in acylcarnitine metabolism in the liver of 3-month old mice treated with PBS or CL-316,243. N = 5-7/group. (C) Expression of hepatic HNF4α targets in 3 month-old mice treated with PBS or CL-316,243. N = 5-7/group. All transcripts were normalized to RPL13, statistical analysis completed used student t-test. Data are presented as mean ± SEM. *p ≤ 0.05, **p ≤ 0.01, ***p≤0.001. See also Figure S3.

### HNF4α regulates acylcarnitine metabolism in the liver of cold exposed mice

The observed increase in hepatic gene expression of enzymes involved in acylcarnitine metabolism led us to test a direct role for HNF4α in the liver, particularly with prior evidence that the nuclear receptors HNF4α and PPARα are known regulators of Cpt1 and Cpt2 expression (Gutgesell et al., 2009; Hayhurst et al., 2001; Louet et al., 2002; Martinez-Jimenez et al., 2010; Song et al., 2010). To determine whether the cold induced changes in hepatic gene expression were due to HNF4α, we generated mice lacking HNF4α in hepatocytes. HNF4α^f/f^ mice received either a control adeno-associated virus 8 (AAV8)-thyroid hormone binding globin (TBG)-eGFP or AAV8-TBG-eGFP-Cre one week prior to cold challenge, leading to selective deletion of HNF4α in hepatocytes (Figure 4A, 4B). After 5 hours of cold exposure, loss of HNF4α led to impaired induction in Cpt1b, Octn2, and CrAT, Apoa4, and PGC-1α when compared to AAV8-TBG-eGFP controls (Figure 4B), while basal levels of Cpt2 and CACT were reduced in HNF4α null mice. These transcriptional changes correlated with functional changes in thermogenesis. HNF4α^f/f^ AAV8-TBG-eGFP-Cre mice were unable to maintain their core body temperatures when challenged with a cold tolerance test, and displayed lower levels of circulating acylcarnitines (Figure 4C and 4E). The loss of HNF4α did not lead to complete disruption of circulating lipids. HNF4α^f/f^ AAV8-TBG-Cre mice exhibited a similar rise in FFAs during cold exposure as the HNF4α^f/f^ AAV8-TBG-eGFP control mice (Figure 4D). To rule out that other sources of energy were not depleted, we measured blood glucose levels, and found an increase in mice lacking HNF4α in hepatocytes (**Figure S4A**). Notably, the expression of thermogenic genes UCP1, Elovl3, Dio2, and brown adipocyte markers, Cidea, Prdm16, and Eva1 were not altered in the brown adipose tissue of hepatocyte-selective HNF4α null mice (**Figure S4B**). Furthermore, gene expression of acylcarnitine transcripts Cpt1b, Octn2, CACT, and CrAT were unchanged in BAT (**Figure S4C**).

**Figure 4:**
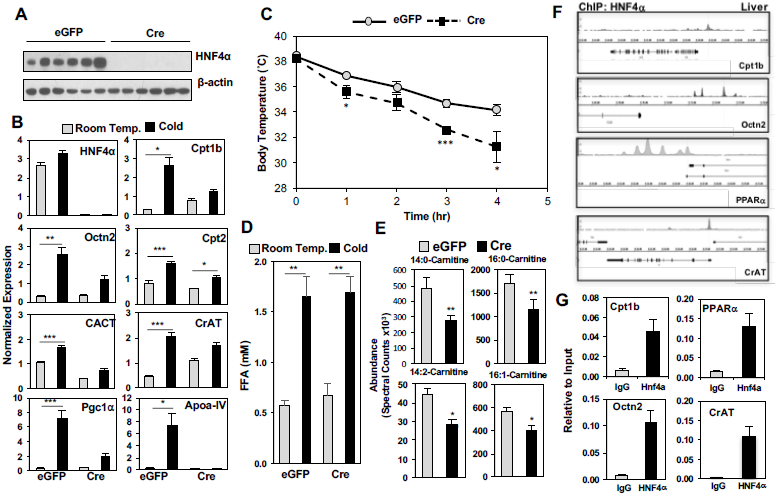
Hepatic HNF4α regulates acylcarnitine metabolism in the liver of cold exposed mice. (A) Western blot analysis of HNF4α and β-actin in the liver of HNF4α^F/F^ mice infected with AAV8-TBG-eGFP (eGFP) or AAV8-TBG-eGFP-Cre (Cre). N = 6/group. (B) Gene expression changes by Real-time PCR analysis of livers from HNF4α^F/F^ mice infected with AAV8-TBG-eGFP (eGFP) or AAV8-TBG-eGFP-Cre (Cre). Mice were challenged with cold or room temperature for 5 hrs. N = 3 for room temperature and N = 6 for cold. (C) Core body temperature after a cold tolerance test at 4°C in HNF4α^F/F^ mice infected with AAV8-TBG-eGFP (eGFP) or AAV8-TBG-eGFP-Cre (Cre). N = 6/group. (D) Free fatty acid (FFA) levels in serum of HNF4α^F/F^ mice infected with AAV8-TBG-eGFP (eGFP) or AAV8-TBG-eGFP-Cre (Cre). Mice were challenged with cold or room temperature for 5 hrs. N = 3 for room temperature and N= 6 for cold. (E) Serum long chain acylcarnitine levels as measured by LC-MS from 5hr cold-exposed HNF4α^F/F^ mice infected with AAV8-TBG-eGFP (eGFP) or AAV8-TBG-eGFP-Cre (Cre). N = 6/group. (F) Hepatic ChIP-seq analysis of HNF4α in proximity to promoters of Cpt1b, Octn2, PPARα, and CrAT. (G) Targeted ChIP-qPCR of HNF4α in the promoters of Cpt1b, Octn2, PPARα, and CrAT in the liver of 3-month old C57BL6J male mice. N = 3 All transcripts were normalized to RPL13. Data are presented as mean ± SEM. *p ≤ 0.05, **p ≤ 0.01, ***p≤0.001. See also Figure S4.

Although Cpt1 and Cpt2 regulation by HNF4α is well established, we also observed reduction in Octn2 and CrAT expression (Hayhurst et al., 2001; Martinez-Jimenez et al., 2010). To determine if these changes in Octn2 and CrAT were due to direct regulation by HNF4α, we interrogated publically available ChIP-Seq data sets to determine whether HNF4α occupies their promoters (Alpern et al., 2014). HNF4α binding peaks were observed in proximity to promoters of PPARα, Cpt1b, Octn2, and CrAT (Figure 4F). These peaks were validated by targeted ChIP-qPCR using livers from C57BL6J mice (Figure 4G). Together these findings suggest that HNF4α is a major regulator of the cold adaptive response in the liver.

### Cold-induced rise in circulating FFAs leads to HNF4α activation in the liver

The observation that liver acylcarnitine metabolism is increased with β3-AR agonist CL-316,243 (Figure 3A) suggests there may be crosstalk between hepatocytes and cells that express the β3-AR. In addition to brown adipocytes, white adipocytes express the β3-AR, which is activated in response to the cold to mobilize FFAs, a potential signal for hepatic HNF4α. To understand the temporal changes in FFA release, we measured FFAs in the serum of mice at 0, 30 minutes, 1 hour, 3 hours, and 5 hours post-cold exposure, and found maximal stimulation within 30 minutes of cold exposure (Figure 5A). In the same experiment, we also measured long chain acylcarnitine levels, and found that 12:1-, 14:0-, 16:0-, and 18:0-carnitine are elevated later, after 3 hours of cold exposure, and continue to rise after 5 hours (Figure 5A, **Figure S5A**). To test whether the rise in FFAs is required for the changes in hepatic gene expression and acylcarnitine metabolism, we treated mice with a single dose of the ATGL inhibitor Atglistatin. Loss of ATGL or Atglistatin treatment prevents cold-induced fatty acid release from adipocytes (Ahmadian et al., 2011; Haemmerle et al., 2006; Heldmaier and Seidl, 1985). Therefore, we treated mice with 200μmol/kg body weight of Atglistatin, preventing the rise in FFAs in response to the cold (Figure 5B), blocking the induction of hepatic HNF4α targets, including ApoA-IV, Cpt1b, CrAT, CACT, Cpt2, and Octn2 (Figure 5C). Blocking the rise in FFAs also prevented the rise in LCACs seen with cold exposure (Figure 5D), and led to an impaired thermogenic response (Figure 5E). Together these findings indicate that fatty acid mobilization is key to activating the transcriptional response in the liver and stimulating the induction in serum acylcarnitine levels.

**Figure 5:**
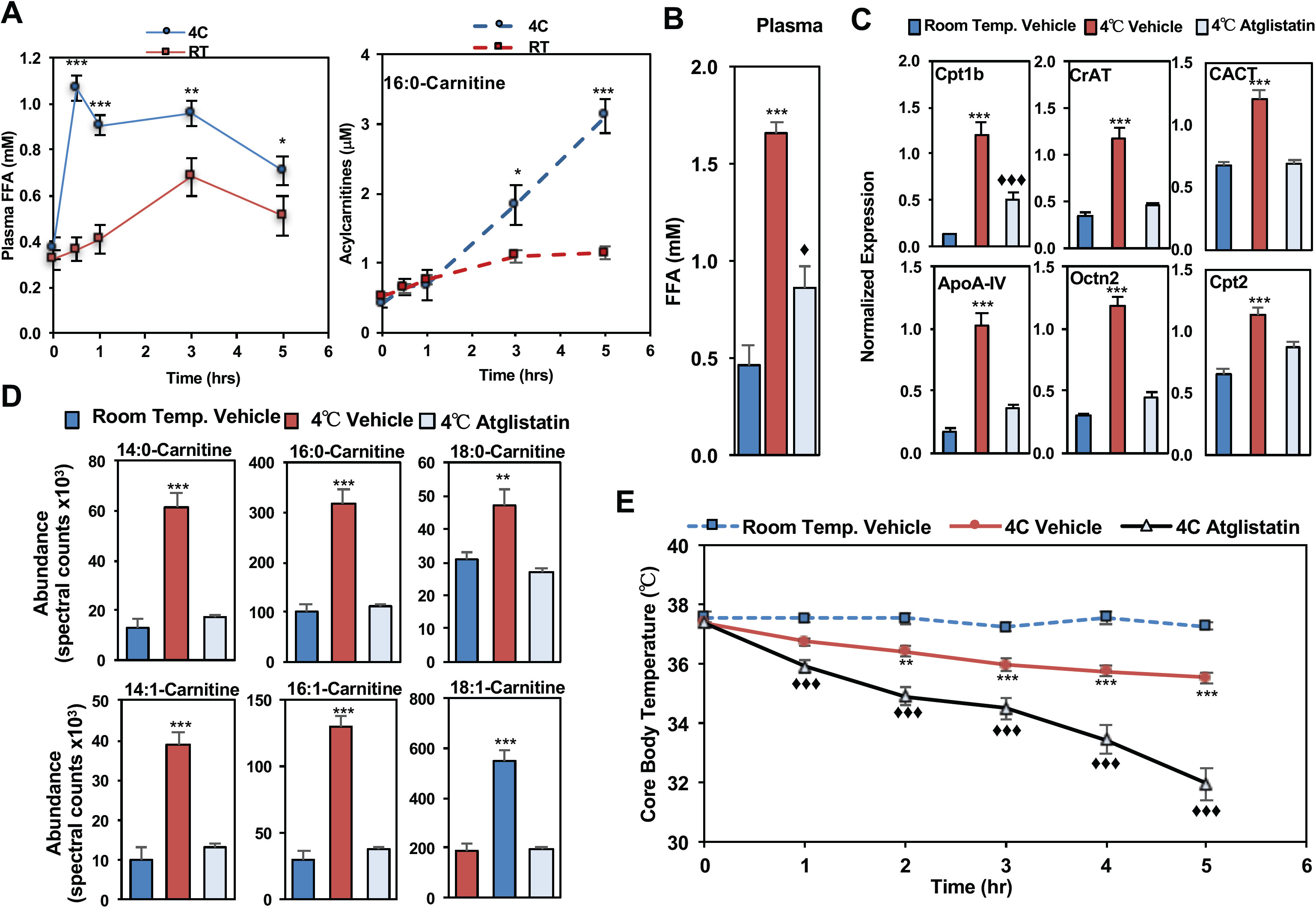
Free fatty acids released from the white adipose tissue are required for increased acylcarnitine production by the liver. (A) Time course of serum free fatty acids (FFA) and palmitoylcarnitine levels of 3 month old male mice exposed to Room Temperature (RT) or Cold (4°C). N = 5/group. (B) Serum free fatty acid (FFA) levels of C57BL6J mice treated with vehicle or Atglistatin at Room Temperature (RT) or Cold (4°C). N = 5/group. (C) Gene expression changes by real-time PCR analysis from livers of C57BL6J mice treated with vehicle or Atglistatin at Room Temperature (RT) or Cold (4°C). N = 5/group. (D) Changes in serum acylcarnitine levels were measured by LC-MS from mice treated with vehicle or Atglistatin at Room Temperature (RT) or Cold (4°C). N = 5/group. (E) Core body temperature measured by rectal probe in mice treated with vehicle or Atglistatin at Room Temperature (RT) or Cold (4°C). N = 5/group. All transcripts were normalized to RPL13. Data are presented as mean ± SEM. For comparison of Room Temp vs. 4°C*p ≤ 0.05, **p ≤ 0.01, ***p≤0.001. For comparison of Room Temp Vehicle to 4°C Atglistatin. ♦ p ≤ 0.05, ♦♦ p ≤ 0.01, ♦♦♦ p≤0.001. See also Figure S5.

### Palmitate treatment in hepatocytes activates expression of acylcarnitine pathway through HNF4α

To test whether free fatty acids could directly stimulate the expression of HNF4α targets, we developed a cell line of HNF4α^f/f^ hepatocytes expressing Rosa 26 LSL-tdTomato, which allows us to test efficiency of Cre recombinase by detection of RFP (Figure 6A). Hepatocytes were infected with adeno-associated virus 2-days prior to treatment with fatty acids. Hepatocytes were incubated with BSA alone or BSA conjugated to palmitate at a concentration of 0.25, 0.5, and 1.0 mM. Palmitate treatment increased the gene expression of HNF4α targets, Octn2, Apoa4, MTTP, and PGC1α in a dose dependent fashion, while HNF4α and cyk18 expression were unaltered. This was abrogated by the loss of HNF4α, as hepatocytes infected with Cre recombinase had a blunted response to palmitate *in vitro* (Figure 6B, **Figure S6A**). These findings suggest that treatment of hepatocytes with free fatty acids is sufficient in stimulating the cold-induced transcriptional response.

**Figure 6:**
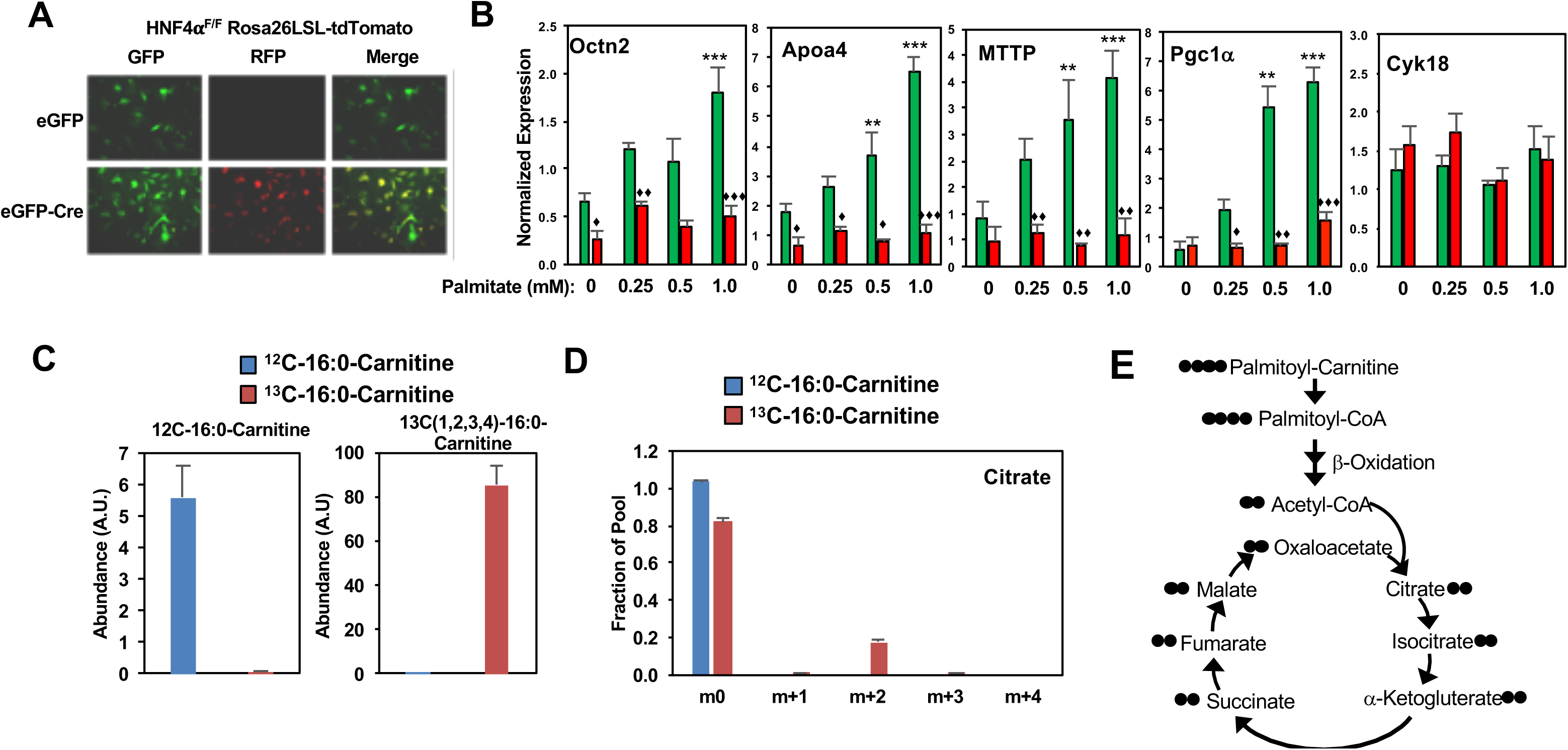
Acylcarnitines are taken up by BAT and metabolized through the TCA cycle. (A) Hepatocytes from HNF4α^F/F^ mice expressing Rosa26-LSL-tdTomato infected with adenoviral CMV-eGFP (eGFP) or CMV-eGFP-Cre (Cre). Cre induced recombination leads to expression of red fluorescent protein (RFP). Hepatocytes were infected for 16hrs two-days prior to harvest. (B) Gene expression changes in hepatocytes measured by real-time PCR after treatment with BSA or BSA/palmitate at 0.25, 0.5, and 1.0 mM for 6 hrs. N = 3/group. (C) Uptake of heavy labeled palmitoylcarnitine as measured by GC-MS in differentiated brown adipocytes incubated with either ^12^C-16:0-Carnitine or ^13^C-(1,2,3,4)-16:0-Carnitine for 6 hours. N = 5/group. (D) Incorporation of ^13^C-(1,2,3,4)-16:0-Carnitine into TCA cycle intermediate m+2 Citrate, and schematic of heavy label entry and incorporation into the TCA cycle. N = 5/group. All transcripts were normalized to RPL13. Data are presented as mean ± SEM. For comparison of HNF4α^F/F^ eGFP between BSA control (0) and palmitate treatment are shown as *p ≤ 0.05, **p ≤ 0.01, ***p≤0.001. N = 5. For comparison between eGFP and Cre infected cells of the same treatment groups ♦ p ≤ 0.05, ♦♦ p ≤ 0.01, ♦♦♦ p≤0.001 See also Figure S6.

### Circulating acylcarnitines provide a fuel source for BAT thermogenesis

The metabolic fate of circulating acylcarnitines is unknown. Using isotopic labeling, we tested whether brown adipocytes metabolize LCACs by incubating differentiated brown adipocytes with 100 μM ^12^C-palmitoylcarnitine or heavy-labeled ^13^C-1,2,3,4-palmitoylcarnitine, and measured the incorporation into the TCA pool. We found that cells take up palmitoylcarnitine (Figure 6C), as supported by the abundance of the m+4 isotopomer of palmitoylcarnitine. To test whether ^13^C-1,2,3,4-palmitoylcarnitine was metabolized, we measured the incorporation of ^13^C into the TCA intermediate Citrate, and found that the m+2 isotopomer could be detected (Figure 6D). Together these findings indicate that palmitoylcarnitine is taken up by brown adipocytes and metabolized.

### Reversal of cold sensitive phenotype in aged mice is rescued by acylcarnitine administration

As mice age, there is an impairment in thermogenic capacity (**Figure S2**), and as shown in Figure 1D, reduced acylcarnitine levels. Carnitine administration increases the abundance of acylcarnitines in the blood (**Figure S7A**). To test whether the reduction with aging impairs thermogenesis, we administered a single injection of 100mg/kg of carnitine, which led to an increase in acylcarnitine levels during cold exposure (Figure 7A). We found that carnitine administration prevents the hypothermia induced by aging, reversing cold sensitivity in 1-year old mice placed at 4°C (Figure 7B) and 2.5-year old mice placed at 16°C (Figure 7C). When given a single bolus of 100mg carnitine per kg body weight the 2.5-year old mice lost on average of 1.5°C compared to the control group that lost 5°C (Figure 7C). Moreover, infusion of palmitoylcarnitine by tail vein was able to improve thermoregulation at 16°C in 2.5-year-old mice with temperatures only dropping 2°C (Figure 7D). The bolus of carnitine did not alter BAT gene expression in the 2.5-year old mice when we assessed UCP1, Dio2, Elov3, and Eva1 (**Figure S7B**). Together these findings indicate that the drop in acylcarnitines contributes to the cold sensitive phenotype observed with aging.

**Figure 7:**
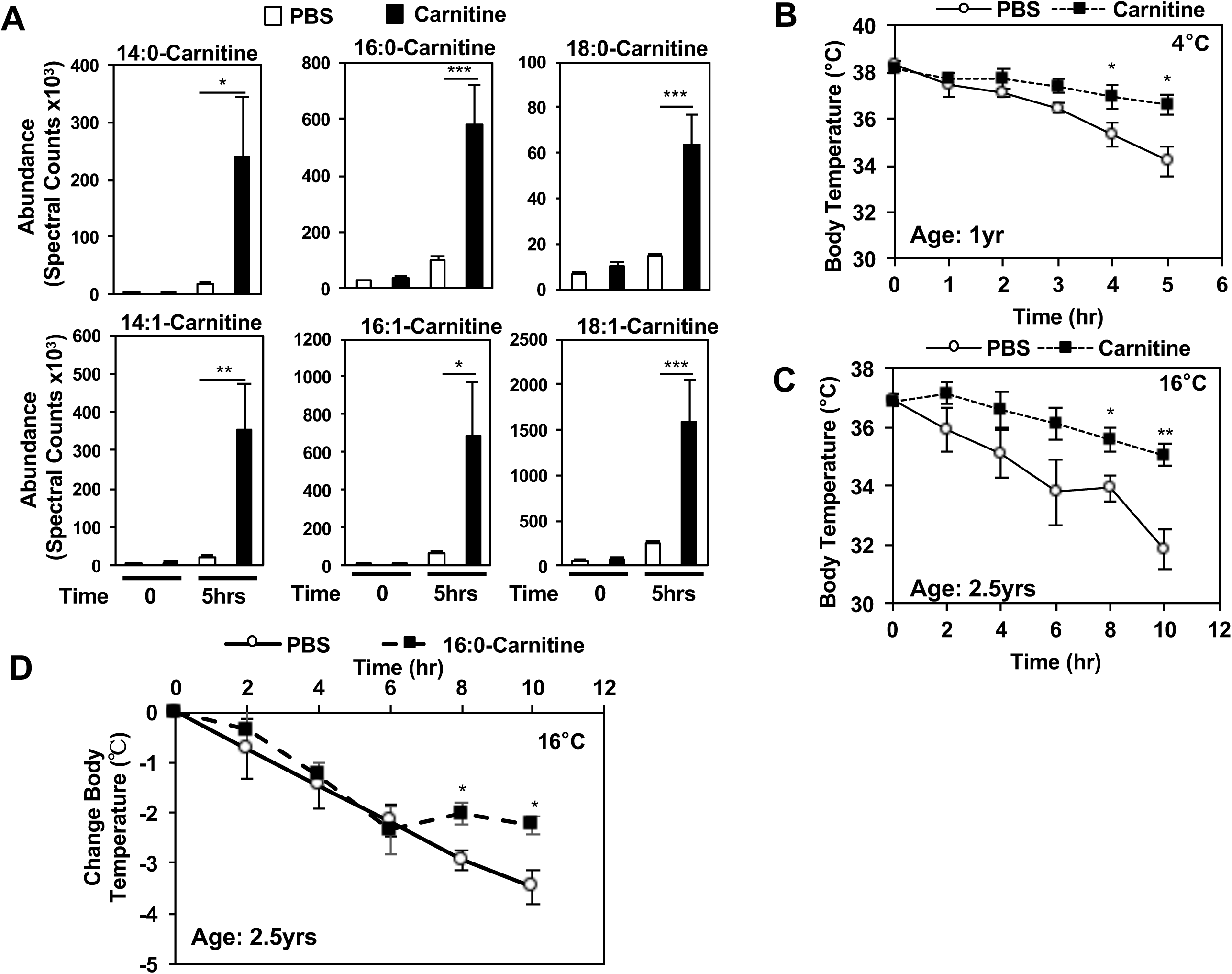
Carnitine treatment stimulates acylcarnitine production and protects against age-induced cold sensitivity. (A) Treatment of 24-month old mice with carnitine (100mg/kg body weight) increases their serum acylcarnitine levels after 5 hours of cold as measured by LC-MS. N = 5. (B) Core body temperature after cold tolerance test at 4°C in 12-month old mice treated with either PBS or 100 mg/kg carnitine. N = 5. (C) Core body temperature after cold tolerance test at 16°C in 24-month old mice treated with either PBS or 100 mg/kg carnitine. N = 5. (D) Change in core body temperature in 24-month old mice treated with PBS or 16:0-Carnitine (100μM). N = 5. Data are presented as mean ± SEM. *p ≤ 0.05, **p ≤ 0.01, ***p≤0.001. N = 5. See also Figure S7.

## Discussion

Lipidomic analysis of plasma lipids during cold exposure led to the identification of acylcarnitines as a cold-induced metabolite. While several acylcarnitine species had previously been detected in the blood in cases of inborn errors of metabolism, exercise, diabetes, and fasting, little was known about their physiologic function. Here, we show that acylcarnitine levels increase in response to the cold and provide a fuel source for BAT thermogenesis, bypassing CPT1, the rate limiting step for fatty acid oxidation. Acylcarnitine production is stimulated through the activation of the nuclear receptor HNF4α, by directly regulating the expression of genes involved in acylcarnitine metabolism. HNF4α activation requires the increase in FFA release from adipose tissue lipolysis, which provides a stimulus for HNF4α activation, and substrate for acylcarnitine synthesis. It has been well documented in rodents and humans that aging impairs thermogenesis, however little is known about the mechanisms at play. Here we found that acylcarnitine levels in response to cold exposure were blunted in older mice. The cold sensitive phenotype observed can be reversed with carnitine administration, a treatment that’s known to stimulate acylcarnitine production. Furthermore, infusing palmitoylcarnitine leads to a similar outcome. In sum, these results suggest a role for the liver in thermogenesis, as well as uncovering a well-orchestrated inter-tissue communication system to upregulate energy mobilization for heat production.

The liver is a focal point for energy mobilization, providing ketones and glucose during fasting, as well as packaging lipid rich lipoproteins for peripheral tissues. Acylcarnitines in the plasma have traditionally been thought of as markers of metabolic stress that increase during fasting, exercise, obesity, inborn errors of metabolism, and now as we observed during cold exposure (Burrage et al., 2016; Mai et al., 2013; McCoin et al., 2015; Schooneman et al., 2013). Although their levels are known to be altered, a functional role for acylcarnitines has yet to be determined. Using the metabolic tracer ^13^C-1,2,3,4-palmitoylcarnitine, we found that acylcarnitines were taken up by brown adipocytes and metabolized, an outcome that was supported by the heavy labeling of the TCA intermediate Citrate. Notably, carnitine supplementation reverses the low levels of circulating acylcarnitines in old mice in response to the cold, and improves their ability to adapt to the cold. Alternatively, acylcarnitines may improve thermoregulation through metabolic flux in the liver, producing heat as a byproduct of acylcarnitine synthesis. Early studies suggest that hepatic thermogenesis is possible, an outcome that was previously measured by calorimetry and shown to contribute to the total body temperature (Baconnier et al., 1979). Further studies are needed to determine the input of hepatic lipid processing in thermoregulation during cold exposure; however, our result that palmitoylcarnitine alone is sufficient to enhance thermogenesis indicates that LCACs are capable of modulating thermogenesis independent of their synthesis, supporting a model that acylcarnitines are a fuel source for thermogenesis.

There is a prevailing view that free fatty acids are the major source of energy for thermogenesis, yet we find that acylcarnitine production is required, suggesting that this long held view is overly simplistic. This is supported by the observation that deletion of hepatic HNF4α leads to reduced acylcarnitine levels, while free fatty acid levels are unchanged, suggesting that free fatty acids are not sufficient for thermogenesis. This finding is surprising, considering the high abundance of free fatty acids that are available for thermogenesis in response to the cold. However, the metabolic program of brown adipocytes is unique compared to other cell types, where its activation leads to enhanced fatty acid oxidation, glucose uptake and utilization, lipogenesis, and lipolysis. This unique metabolic program reflects the dualistic nature of BAT that is capable of storing excess lipids similarly to other adipose depots, and yet shares similar features with skeletal muscle, including a common precursor and high mitochondrial content (Festuccia et al., 2011; Yu et al., 2002). In other tissues fuel selection is regulated by the Randle cycle, a series of inhibitory signals that decrease glucose or fatty acid utilization, and is dependent on their availability (Hue and Taegtmeyer, 2009). In the Randle cycle when both glucose and fatty acids are readily available, the TCA-cycle intermediate citrate, is exported to the cytoplasm to ultimately generate malonyl-CoA, an inhibitor of Cpt1b, the rate limiting step in fatty acid oxidation. Through inhibition of fatty acid oxidation, this allows the preferential use of glucose for energy (McGarry et al., 1991). Although cold exposure is energetically demanding, there is a high abundance of peripheral fuel sources; both circulating glucose, regulated by a surge in glucagon, and FFAs are readily available (Kinoshita et al., 2014; Wu et al., 2006). During acute cold exposure malonyl-CoA levels in BAT rise to levels that would inhibit Cpt1 activity (Saggerson and Carpenter, 1982). These conditions suggest that initial BAT thermogenesis is dependent on glucose uptake, while persistent cold challenge allows for a switch to utilization of the more energetically rich lipids. Therefore, we propose a model where acylcarnitines provide a mechanism for lipid utilization to bypasses the inhibition of Cpt1b, expediting the metabolic switch from glucose to FFA utilization.

The increase in expression of genes involved in acylcarnitine metabolism in response to cold was surprising, as there is little evidence for the liver’s involvement in adaptive thermogenesis. Prior studies on the transcriptional regulation of Cpt1 led our focus to HNF4α, which is primarily thought to be involved in liver development. Analysis of HNF4α occupancy using publically available ChIP-Seq data sets revealed potential binding sites in the promoter region of other genes involved in acylcarnitine metabolism (Louet et al., 2002; Martinez-Jimenez et al., 2010). Notably, HNF4α binding at these sites did not change in response to the cold, an outcome noted by other HNF4α targets (data not shown); instead it is the binding to various coactivators that changes HNF4α activity. HNF4α activity is largely driven by coregulator interactions, where Hes6 is thought to be inhibitory, while interactions through PGC-1α activate transcription (Martinez-Jimenez et al., 2010; Rhee et al., 2006; Rhee et al., 2003). We found that in response to the cold, expression of hepatic PGC-1α was induced, but lost as mice aged. The induction of PGC1α is likely driven by FFAs, which stimulate CREB phosphorylation, and increase expression of PGC1α. These findings fit with our model that HNF4α activation is dependent on the FFA release observed with the cold or activation with a β3-AR agonist CL-316,243 (Collins et al., 2006; Herzig et al., 2001; Schauer and Reusch, 2009). Alternatively FFAs could activate HNF4α directly through its ligand binding domain, although binding of HNF4α to its purported lipid ligand has been shown to occur during assembly, while others have shown linoleic acid regulates HNF4α activity (Dhe-Paganon et al., 2002; Yuan et al., 2009). Our studies indicate that HNF4α is required for the induction of PGC1α, Octn2, MTTP, and Apoa4 in response to palmitate in hepatocytes. Further studies are needed to elucidate the mechanism through which cold exposure increase HNF4α activity as well as other regulatory elements controlling this cold response in the liver. Although the induction of genes involved in acylcarnitine metabolism are abrogated in HNF4α knockout livers, they are not completely ablated. This suggests that other transcriptional regulators are involved in mediating the cold response observed with Octn2, Cpt2, CACT, and CrAT. Together our findings support a model where FFA release during the cold, stimulates a transcriptional program driven by HNF4α, ultimately leading to acylcarnitine synthesis and release into the plasma. This model explains why the β3-AR agonist CL-316,243 stimulates acylcarnitine levels in the blood, despite the lack of β3-AR expression in hepatocytes.

The rise in acylcarnitines is dependent on the increase in circulating FFAs, and inhibition of triglyceride lipolysis by a single dose of ATGL inhibitor, Atglistatin, blocked the increase in circulating acylcarnitines in response to cold exposure. Atglistatin treatment also led to cold intolerance in mice, however, this could reflect the dependence of FFAs as a fuel source for thermogenesis, or the failure to activate of Ucp1 in BAT by inhibiting lipolysis (Fedorenko et al., 2012; Wu et al., 2006). Although Atglistatin treatment inhibits ATGL in multiple tissues, others have shown that ATGL knockout mice are sensitive to hypothermia, and that loss of ATGL in the adipocytes is sufficient to decrease β3-AR agonist stimulation of FFA release (Ahmadian et al., 2011; Haemmerle et al., 2006). Interestingly, Atglistatin treatment also inhibited the increased expression of hepatic acylcarnitine transcripts, suggesting that the cold induced FFAs are not only acting as a substrate for acylcarnitine synthesis but also activate the transcriptional regulatory program.

Another model that highlights the importance of acylcarnitine availability in thermogenesis is found in aged mice. As mice and humans age they have decreased BAT activity, are prone to hypothermia, and have decreased circulating acylcarnitines despite an abundance of peripheral fuel sources including FFAs (Houtkooper et al., 2011; Sellayah and Sikder, 2014; Yoneshiro et al., 2011). We found that older mice had decreased levels of LCACs and a single carnitine bolus was able to restore LCAC levels and improve thermogenic capacity. The ability of carnitine supplementation to enhance the thermogenic capacity of older mice could reflect an enhanced shivering or non-shivering thermogenesis. As mice age they experience sarcopenia as well as loss of BAT, and previously carnitine supplementation has been shown to increase muscle pyruvate dehydrogenase activity, mitochondrial function and glucose homeostasis in mice and humans (Burrage et al., 2016; Muoio et al., 2012; Seiler et al., 2015). The mechanism by which carnitine supplementation increases acylcarnitine levels in the context of our aging studies where total carnitine levels are not limited remains elusive. Although others have seen a decrease in total carnitine levels with aging and overnutrition, we did not observe this decrease in the 24 month old mice (Noland et al., 2009). However, the lower abundance of all chain lengths of acylcarnitines suggests a decrease in the carnitine pool that the carnitine supplementation may ameliorate, and alludes to an unknown regulatory system controlling carnitine flux. More work is needed to understand the mechanism by which carnitine supplementation improves thermoregulation, however, its ability to amend hypothermia in aged mice may offer therapeutic potential in the treatment of cold sensitivity in elderly populations.

Thermogenesis is critical for survival, and has allowed mammals to survive the cold. Our studies aimed to develop a better understanding of the metabolic changes that take place with cold exposure. Lastly, this paper presents several seminal discoveries that will have broader impacts on the fields of BAT thermogenesis and lipid homeostasis. We observed an increase in circulating acylcarnitines with β3AR activation, and propose that these acylcarnitines are produced by the liver and taken up by the BAT as a fuel for thermogenesis. These findings also represent a novel role for the liver and activation of HNF4α in acute cold exposure. This broadens the understanding of the liver as a metabolic hub that processes fuel sources for other tissues including hepatic gluconeogenesis, lipoprotein synthesis, ketogenesis, and now acylcarnitine production. In sum, these studies demonstrate the importance of peripheral energy sources in heat production by brown adipocytes and discovered an inter-tissue communication system that regulates thermogenesis.

## Experimental Procedures

### Animal Studies

C57BL6J male mice aged to 3 months were purchased from Jackson laboratories, older mice were provided by the National Institute on Aging. HNF4α^f/f^ mice were previously described (Hayhurst et al., 2001). HNF4α^f/f^ mice were aged to 12weeks, one week prior to experimental harvest mice received intraperitoneal injection of adeno-associated virus 8 (AAV8) containing Cre recombinase regulated by the thyroid hormone binding globin (AAV8-TBG-Cre) or a control green fluorescent protein regulated by the same promoter (AAV8-TBG-eGFP) (University of Pennsylvania Vector Core in Philadelphia). Mice received a single injection of 400μL of AAV8-TBG-eGFP or AAV8-TBG-Cre (titer 10^12^gnome copies/mL).

Administration of CL-316,243 (1mg/kg body weight; Cayman) and carnitine (100mg/kg body weight; Sigma) was performed by intraperitoneal injection, while palmitoyl-carnitine (100μM; Sigma) and palmitate conjugated to BSA (100μM; Sigma) was injected via tail vein. Sterile PBS pH 7.5 was used as a vehicle control and administered in the same manner as the experimental group. Atglistatin (200umol/kg body weight; Caymen) was diluted in corn oil and provided by oral gavage, a comparable volume of corn oil was administered by gavage in the control mice (Mayer et al., 2013). Prior to cold exposure mice were singly housed in a cage with no food, no bedding, but ready access to water. Mice were placed in either 24°C (room temperature), 16°C (activation of non-shivering thermogenesis), or 4°C (cold exposure) for 5 hours. Their core body temperature was monitored hourly by rectal probe.

### Cell Culture

Brown preadipocytes were isolated from C57BL6J mice as previously described and were immortalized through retroviral expression of SV40 Large T-antigen using neomycin for selection (Rodriguez-Cuenca et al., 2007). Cells were plated in DMEM containing 10% FBS (RMBI), 20nM insulin (Sigma), and 1nM T3 (Sigma). Upon complete confluence cells were stimulated for differentiation by DMEM containing 10% FBS, 20nM insulin, 1nM T3, 0.5mM isobutylmethylxanthine (Sigma), 0.5μM dexamethasone (Sigma), 0.125mM indomethacin (Sigma), and 1μM Rosiglitazone (Cayman). After 2 days differentiation media was removed and cells were maintained in DMEM 10% FBS, 20nM insulin, 1nM T3, and 1μM Rosiglitazone. Six days after the addition of differentiation media cells were washed 3 times with PBS, then treated with 100μM ^12^C-palmitoylcarnitine (Sigma) or 100μM ^13^C-1,2,3,4-palmitoylcarnitine (Isotec) in Krebs Ringer Buffer (135mM NaCl, 5mM KCl, 1mM MgSO_4,_ .4mM K_2_HPO_4,_ 5.5mM Glucose, 20mM HEPES, 1mM CaCl_2_ pH 7.4). After 6 hours of palmitoylcarnitine treatment cells were washed 3 times with PBS and then collected by cell scrapper for metabolic tracer analysis.

Primary hepatocytes were isolated as previously described (Severgnini et al., 2012) from HNF4α^f/f^ Rosa26LSL-tdTomato a kind gift from the laboratory of Eric Snyder at the University of Utah. Briefly, the livers were perfused with sterile PBS through p10 tubing inserted into the vena cava, the visceral vena cava was cut to allow flow and then clamped for pressure to ensure complete perfusion. Liver was then perfused with HBSS, followed by DMEM with .15% Collagenase (Sigma). The cells were passed through a 100μM cell strainer and then pelleted at 50 g for 1 minute, the pellet was washed 3 times with DMEM and then plated on a tissue culture plate with 0.1% rat tail collagen (Sigma). Cells were maintained in hepatic cell culture media (ThermoFisher), and assessed for hepatocyte purity by RT-PCR of albumin, cyk18, and transthyretin. To generate HNF4α control and knock out cells, hepatocytes were infected with adenovirus expressing either CMV-eGFP (GFP) or CMV-eGFP-Cre (CRE) (ViraQuest). 48 hours post infection cells were treated with various levels of BSA conjugated palmitate for 6 hours, washed 3 times with PBS, and then RNA was extracted.

### Lipid Measurements

Lipids were extracted from serum (∼40 μL) aliquots then combined with 225 μL ice-cold MeOH containing internal standards (Avanti Lipids, LM-1601 (19.82 μM, -1102 (14.52 μM) and -1002 (12.54 μM); 10 μL each / sample) and vortexed for 10 s. 750 μL of ice-cold MTBE (methyl tert-butyl ether) was added, vortexed for 10 s, and 200 μL of water is added to induce phase separation. The sample was then vortexed for 20 s followed by centrifugation at 14,000 g for 2 min at 4 °C. The upper phase (750 μL) was collected and evaporated to dryness under vacuum. Samples were reconstituted in 25 μL ACN:H_2_O:IPA (1:1:2 *v/v*) + 0.1% formic acid for analysis. A pooled QC sample was prepared by 5 μL aliquots from each sample.

Lipid extracts were separated on an Acquity UPLC CSH C18 1.7 μm 2.1 x 100 mm column maintained at 60 °C connected to an Agilent HiP 1290 Sampler, Agilent 1290 Infinity pump, equipped with an Agilent 1290 Flex Cube and Agilent 6520 Accurate Mass Q-TOF dual ESI mass spectrometer. For positive more, the source gas temperature was set to 350 °C, with a gas flow of 11.1 (L/min) and a nebulizer pressure of 24 psig. VCap voltage is set at 5000 V, fragmentor at 250 V, skimmer at 74.4 V and Octopole RF peak at 750 V. VCap voltage is set at 5000 V, fragmentor at 100 V, skimmer at 75 V and Octopole RF peak at 750 V. Reference masses in positive mode (m/z 121.0509 and 922.0098) were infused with nebulizer pressure at 2 psig. Samples were analyzed in a randomized order acquiring with the scan range between m/z 100 – 1700. Mobile phase A consists of ACN:water (60:40 v/v) in 10 mM ammonium formate and 0.1% formic acid, and mobile phase B consists of IPA:water (90:10 v/v) in 10 mM ammonium formate and 0.1% formic acid. The chromatography gradient for both positive and negative modes starts at 15% mobile phase B increasing to 30% B over 4 min, it then increases to 52% B from 4-5 min, then increases to 82% B from 5-22 min, then increases to 95% B from 22-23 min, and then increases to 99% B from 23-27 min. From 27-38 min it’s held at 99%B, then returned to 15% B from 38-38.2 min and was held there from 38.2-44 min. Flow is 0.3 mL/min throughout. Tandem mass spectrometry was conducted using the same LC gradient and at collision energies of 10 V, 20 V and 40 V. Injection volume was 3 μL.

Results from LC-MS experiments were collected using Agilent Mass Hunter Workstation and analyzed using the software package Agilent Mass Hunter Qual B.05.00. Using the Find By Formula (FBF) algorithm, MS/MS fragmentation and a Lipids PCLD database, possible assignments were generated then individually inspected. Compounds were also checked against the blank process sample to remove any artifacts. Once a list of confident assignments was made, a Mass Hunter Quant method was generated and the software program Mass Hunter Quant is used to analyze and integrate each compound. Lipids are normalized to LM-1601 response (PC(17:1(10Z)/0:0)). Assessment of lipidomics t-test p value, fold change, and generation of heat map were performed in MetaboAnalyst 3.0 (Xia et al., 2015). Comparison of the differentially abundant plasma lipids from 3 month old and 24 month old mice in Room Temp or Cold was performed by significance analysis of microarrays (SAM) (Tusher et al., 2001). A tuning parameter, delta of 0.4, optimized the cutoff for significance with the estimation of false discovery rate (FDR) threshold q-value of 0.05. Volcano plot creation and SAM was performed using R (version 3.1.3). Serum free fatty acids were quantified by colorimetric kit according to the manufacturer’s instruction (Sigma).

### Metabolic Tracer Analysis

Brown adipose tissue was weighed in bead mill tubes containing 1.4 mm ceramic beads (Qiagen, Carlsbad, CA). Cold methanol containing *d*4-succinate as an internal standard was added to give a final methanol concentration of 80%. An Omni Bead Ruptor (Omni-Inc, Kennesaw, GA) was employed at 6.45 MHz for 30 seconds to disrupt the cells. The supernatant was transferred to fresh microfuge tubes and protein was precipitated by incubation at -20° C for 30 minutes. The extract was clarified by centrifugation at 20,000 x *g* followed by transfer to new fresh microfuge tubes and solvent removed *en vacuo*.

All GC-MS analysis was performed with a Waters GCT Premier mass spectrometer fitted with an Agilent 6890 gas chromatograph and a Gerstel MPS2 auto sampler. Dried samples were suspended in 40 μL of a 40 mg/mL O-methoxylamine hydrochloride (MOX) in pyridine and incubated for one hour at 30° C. To auto sampler vials was added 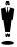 L of this solution. 40 μL of N-methyl-N-trimethylsilyltrifluoracetamide (MSTFA) was added automatically via the auto sampler and incubated for 60 minutes at 37° C with shaking. After incubation 3 μL of a fatty acid methyl ester standard (FAMES) solution was added via the auto sampler then 1 μL of the prepared sample was injected to the gas chromatograph inlet in the split mode with the inlet temperature held at 250° C. A 10:1 split ratio was used for analysis. The gas chromatograph had an initial temperature of 95° C for one minute followed by a 40° C/min ramp to 110° C and a hold time of 2 minutes. This was followed by a second 5° C/min ramp to 250° C, a third ramp to 350°C, then a final hold time of 3 minutes. A 30 m Phenomex ZB5-5 MSi column with a 5 m long guard column was employed for chromatographic separation. Helium was used as the carrier gas at 1 mL/min. Data was collected using MassLynx 4.1 software (Waters). Each isotope for the targeted metabolites were identified and their peak area was recorded using QuanLynx. All the extracted data was then corrected mathematically to account for natural abundance isotopes, and finally showed as the fraction of the total pool of citrate (Katajamaa and Orešič, 2005).

### Gene Expression

RNA was isolated from liver, skeletal muscle, or BAT using Trizol reagent (Invitrogen), samples were homogenized with a TissueLyzer II (Qiagen). Reverse transcription was performed with SuperScript VILO Master Mix (Thermofisher). Quantification of gene expression was performed with KAPA SYBR FAST qPCR 2x Master Mix Rox Low (Kapabiosystems) on an Applied Biosystems QuantStuio 6 Flex Real-Time PCR System, 384-well. Analysis was performed by a calculated relative expression extrapolated from a standard curve for each primer pair that was then normalized to expression of the housekeeping gene RPL13. Primer pairs were designed with Universal Probe Library (Roche) or qPrimer Depot (mouseprimerdepot.nci.nih.gov), a list of primer pairs is included in Table S1. Two-way ANOVA and Tukey post-hoc analysis was performed in Prism 6.

### Protein Analysis

Liver tissue was homogenized by glass dounce in RIPA buffer (Boston Bioproducts) with added protease inhibitor (Roche). Tissue samples were spun at 12,000 x g for 10 minutes at 4°C and the supernatant was extracted. Protein quantification was performed by BCA assay (Thermofisher), diluted in Laemmli loading buffer (BioRad), heated at 70°C for 20 min, and run on a standard 10% acrylamide gel. Protein was transferred to Amersham Protran .45μM Nitrocellulose (GE healthcare) and blotted for Cpt1b (15703, Abcam), Octn2 (16331-1-AP, ProteinTech), Apoa-IV (5700S, Cell Signaling), HNF4α (PP-H1415-00, R & D Systems), or β-Actin (3700S, Cell Signaling).

### Chromatin Immunoprecipitation

Chromatin was prepared from snap frozen livers. Livers were minced in 1% formaldehyde in PBS and incubated for 10 minutes at room temperature. Followed by quenching with 125 mM glycine for 10 minutes at room temperature. Livers were then homogenized in tissue lyser II according to manufacturer’s instructions. Samples were centrifuged at 400g for 5 min and resuspended in chip lysis buffer (25mM Tris pH8, 2mM EDTA, 150mM NaCl, 1% Triton X-100, 0.1% SDS) and sonicated in Diagenode pico bath sonicator for 33 cycles, 40 seconds on 40 seconds off. Chromatin was Immunoprecipitated with antibodies against IgG (PP64 EMD Millipore), HNF4α (PP-H1415-00, R & D Systems) overnight at 4°C in the presence of Dynabeads protein A (10001D, Invitrogen). DNA was purified with ChIP DNA Clean & Concentrator Kit (11-379, Genesee Scientific) and quantified by real-time PCR (ABI QuantStudio Flex6) using Syber green (KK4621, Kapa Biosystems). Occupancy was quantified using a standard curve and normalized to input DNA. Primers are listed in table S1.

## AUTHOR CONTRIBUTIONS

J.S., G.G., A.M., M.P., S.L., P.J.S., I.H., J.C. conducted experiments, J.S., A.J.D., U.A., N.L., J.R., C.J.V. designed experiments, and J.S. and CJV wrote paper. All authors contributed to data analysis.

## ACKNOWLEDGEMENTS

The authors are grateful to members of the Diabetes and Metabolism Center and the Biochemistry Department at the University of Utah for useful discussion and feedback. Lipidomics and metabolic tracer analysis was performed at the Metabolomics Core Facility at the University of Utah. This study was supported by NIDDK KO1DK097285, NIDDK RO3DK103089, NIDDK RO1DK103930, Margolis Research Foundation, NIDDK DRC, T32DK091317, and S10 OD016232-01. The content is solely the responsibility of the authors and does not necessarily represent the official views of the National Institutes of Health

